# Bacteriophage-driven DNA inversions shape bacterial functionality and long-term co-existence in *Bacteroides fragilis*

**DOI:** 10.1101/2025.01.27.635102

**Authors:** Shaqed Carasso, Roni Keshet-David, Jia Zhang, Haitham Hajjo, Dana Kadosh-Kariti, Tal Gefen, Naama Geva-Zatorsky

**Author notes:** These authors contributed equally (Shaqed Carasso, Roni Keshet-David).

## Abstract

Bacterial genomic DNA inversions, which govern molecular phase-variations, provide the bacteria with functional plasticity and phenotypic diversity. These targeted rearrangements enable bacteria to respond to environmental challenges, such as bacteriophage predation, evading immune detection or gut colonization.

This study investigated the short- and long-term effects of the lytic phage Barc2635 on the functional plasticity of *Bacteroides fragilis*, a gut commensal. Germ-free mice were colonized with *B. fragilis* and exposed to Barc2635 to identify genomic alterations driving phenotypic changes.

Phage exposure triggered dynamic and prolonged bacterial responses, including significant shifts in phase-variable regions (PVRs), particularly in promoter orientations of polysaccharide biosynthesis loci. These shifts coincided with increased entropy in PVR inversion ratios, reflecting heightened genomic variability. In contrast, *B. fragilis* in control mice exhibited stable genomic configurations after gut adaptation. The phase-variable Type 1 restriction-modification system, which affects broad gene expression patterns, showed variability in both groups. However, phage-exposed bacteria displayed more restrained variability, suggesting phage-derived selection pressures. Our findings reveal that *B. fragilis* employs DNA inversions to adapt rapidly to phage exposure and colonization, leveraging genomic variability for resilience. This study emphasizes gut bacterial genomic and phenotypic plasticity upon exposure to the mammalian host and to bacteriophages.

## Introduction

The human gut microbiome is a complex and dynamic ecosystem where bacteria must adapt quickly to survive. Among the factors gut bacteria need to cope with are the physiological state of the host neighboring bacteria, and an immense repertoire of bacteriophages^1–3^. Reversible DNA inversions allow gut bacteria to adapt to these ever-changing conditions by altering bacterial functionalities^4–7^. Bacterial DNA inversions are often mediated by inverted repeats, short, palindromic DNA sequences that occur on opposite strands of the DNA. These sequences form secondary structures, such as hairpins, which are recognized by site-specific recombinases^8,9^. The recombinases bind to the inverted repeats, leading to the inversion of the segment between the repeats. These inversions frequently involve promoter regions that control the initiation of transcription, functioning as “ON/OFF” switches^10,11^. Additionally, they can occur within or adjacent to coding regions, shuffling active genes within a locus or creating new genes through the recombination of adjacent partial genes^12–17^.

The Bacteroidales order, abundantly found in the human gut microbiome, has a high prevalence of genomic DNA inversions^11^. Of specific interest is *Bacteroides fragilis*. This bacterial species can express eight distinct capsular polysaccharides (PS), labeled PSA to PSH, seven of which are controlled through DNA inversions at their promoter regions, toggling between “ON” and “OFF” expression states^10,18^. Studies have shown that *B. fragilis’* polysaccharide A (PSA) modulates the host immune system by inducing regulatory T-cells (Tregs) to secrete the anti-inflammatory cytokine interleukin (IL)-10^19,20^. Moreover, PSA has been shown to confer protection against experimental colitis^19,21–23^ and is regarded as an anti-inflammatory polysaccharide.

Our previous work characterized a phase-variable Type 1 restriction-modification (RM) system in *B. fragilis*, which is sensitive to host-derived cues^13^. Type 1 RM systems are bacterial defense mechanisms that consist of a restriction enzyme and a methyltransferase, which work together to protect the bacteria from foreign DNA^24^. In *B. fragilis*, DNA inversions within the Type 1 RM system allow for the expression of eight different specificity proteins, each directing a unique DNA methylation pattern on the bacterial chromosome. These methylation patterns, in turn, regulate distinct transcriptional programs, contributing to the phenotypic diversity and adaptability of the bacteria^13^. The variation in these transcriptional programs can influence the expression of surface structures such as capsular PSs. As a result, these changes in gene expression could potentially alter the immunomodulatory capabilities of *B. fragilis*, allowing it to better adapt to the host environment and interact with the immune system.

In a different study, we recently showed that gut *Bacteroidales* exhibit distinct DNA inversion patterns during gut inflammation, observed both in mouse models and patients with inflammatory bowel disease (IBD)^25^. These inversion patterns are influenced by the host’s inflammatory state, with an increased abundance of *B. fragilis*-associated bacteriophages detected in IBD patients exhibiting the “OFF” orientation of the PSA promoter. Furthermore, exposure of *B. fragilis* to the isolated lytic bacteriophage Barc2635 resulted in a higher frequency of the PSA promoter “OFF” orientation, which altered the host immune-modulatory effects of *B. fragilis* in mice^25^.

Several studies have shown that the interactions between *Bacteroidales* and their associated bacteriophages were shown to play a critical role in altering bacterial behavior and functionality. The surface molecules displayed on *Bacteroides* species can influence their susceptibility to specific phages^14,26^. In *Bacteroides intestinalis*, the ability to modify capsular polysaccharides supports stable coexistence with CrAss-like phages^27^. Phages can also directly impact bacterial functions, as observed in *Bacteroides vulgatus*, where phage infection alters bile salt hydrolase activity through genome integration^28^.

In this study, we aimed to explore the long-term effects of the lytic *Bacteroides*-associated bacteriophage Barc2635 on the functional plasticity of the gut commensal *B. fragilis*. By colonizing germ-free (GF) mice with *B. fragilis* and introducing the phage Barc2635, we sought to observe genomic alterations in the bacteria that can result in phenotypic changes and affect bacterial functional interactions with the host.

## Results

### Barc2635 and *B. fragilis* exhibit long-term persistence in the mammalian gut

To assess the longitudinal effects of Barc2635 on *B. fragilis*, we monocolonized germ-free (GF) mice with *B. fragilis* NCTC 9343 either in the presence or absence of bacteriophage Barc2635. Fecal samples were collected at shorter intervals [see methods, Figure 1A] during the first week to monitor the immediate adaptation of bacterial populations to the phage, and then at weekly intervals [Figure 1A]. In the group of mice colonized with *B. fragilis* alone, bacterial levels fluctuated between 10^11^ and 10^13^ CFU/g feces during the first 200 hours, then stabilized around 10^10^ to 10^12^ CFU/g feces for the remainder of the experiment [Figure 1B]. In the group co-colonized with *B. fragilis* and Barc2635, we observed a dynamic interplay between bacterial and phage populations. Bacterial levels in this group fluctuated throughout the experiment, exhibiting lower CFUs counts than in the control group for most of the timepoints, ranging between 10^10^ and 10^12^ CFU/g feces [Figure 1B]. Phage levels showed a rapid increase after the first week, peaking at around 10^13^ PFU/g feces. These high phage levels (10^12^ to 10^13^ PFU/g) were maintained for several hundred hours, followed by a gradual decline towards the end of the experiment, stabilizing at approximately 10^12^ PFU/g feces [Figure 1B].

**Figure 1:**
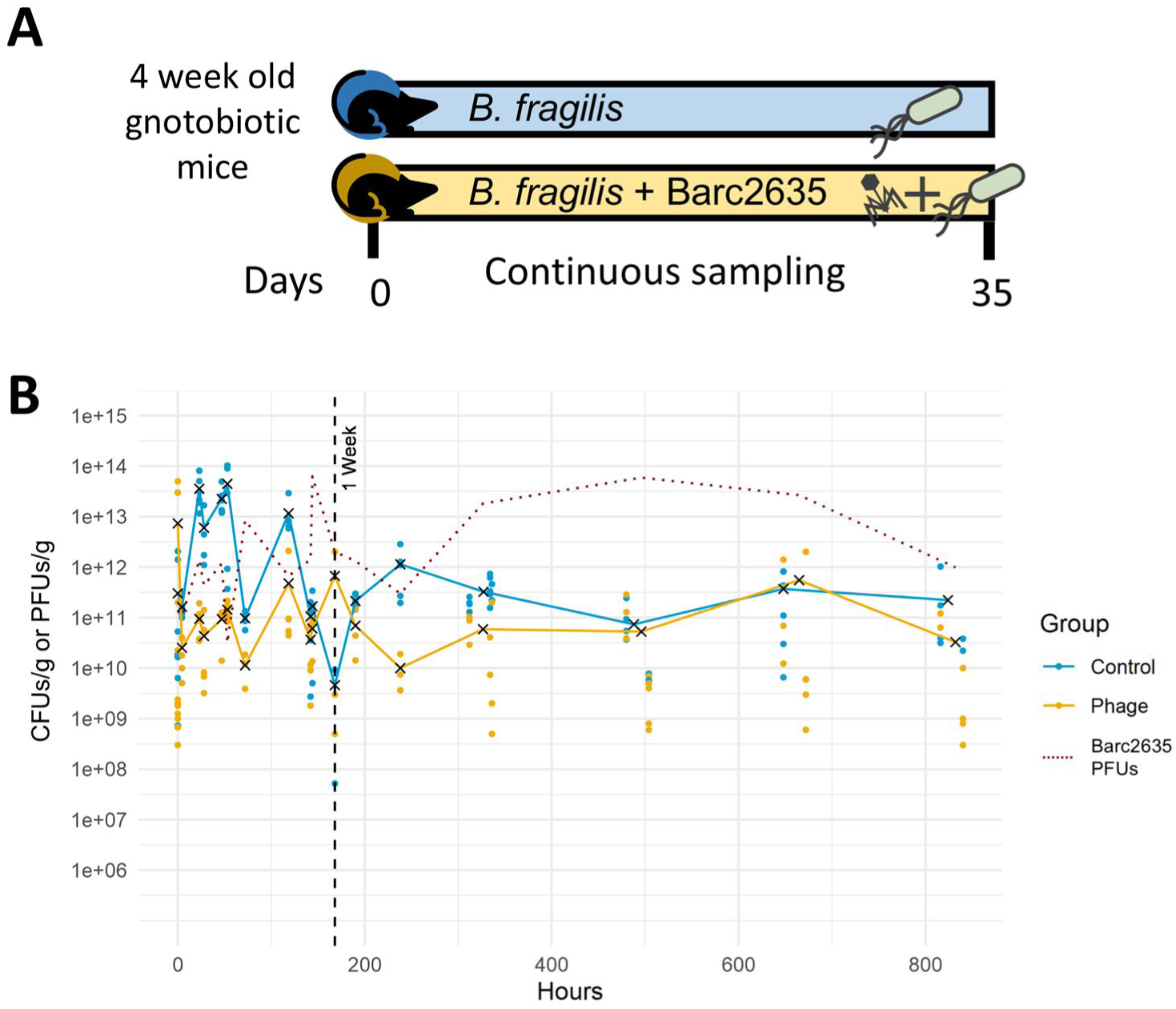
Long-term coexistence of *B. fragilis* and bacteriophage *in vivo*. **(A)** Experimental design of culturing *B. fragilis* and bacteriophage together in GF mice. Four week old mice received oral gavage of *B. fragilis* (Control; blue), or *B. fragilis* + Bacteriophage Barc2635 (Phage; yellow) at a multiplicity of infection (MOI) = 10, and their feces were sampled continuously. **(B)** CFUs of *B. fragilis* in fecal samples of mice monocolonized with *B. fragilis* (blue) or colonized with *B. fragilis* + Barc2635 (yellow) in hours after experiment initiation. Dots represent each fecal sample, the ‘X’ represents the mean of CFUs/g feces. PFUs of Barc2635 in fecal samples of mice colonized with *B. fragilis* + Barc2635 are represented by a dashed red line. The vertical black dashed line represents the one-week timepoint (168 hours).

### B. fragilis adaptation to the mammalian gut and bacteriophage through invertible DNA regions

To investigate the role of invertible DNA regions in *B. fragilis* adaptation, we first conducted an initial experiment using PhaseFinder^11^ on metagenomic sequencing results from the bacteriophage-exposed group. In this analysis, we compared the orientation of phase-variable regions between day 7 and day 35 after introducing the phage and bacteria into GF mice. Based on these results, we selected 18 DNA invertible regions (including the 7 PS loci and 11 additional phase-variable sites) that exhibited either higher ratios of reverse-oriented reads or a large change in orientation between the two timepoints [Table S1]. These regions were then monitored with higher sampling frequency in subsequent experiments, both in the presence and absence of bacteriophage Barc2635.

To evaluate the extent of DNA inversions in both models, we first calculated the entropy of the DNA inversions in the 18 areas we focused on. We valued the entropy as a measure of the variability of the DNA inversions: a measure of both amount and diversity in genomic sites of DNA inversions in both models (i.e. *B. fragilis* with and without bacteriophage exposure). We observed opposite trends of the entropy of *B. fragilis* DNA inversion ratios with and without exposure to phage. In phage-exposed mice, the bacterial DNA inversion entropy increased dramatically during the first 48h and stayed at the higher level, compared to the initial state, and compared to the control group monocolonized with *B. fragilis*, without exposure to bacteriophages, indicating high levels of disorder. On the other hand, in the control group where *B. fragilis* was not exposed to bacteriophage, the entropy decreased after the bacterial inoculation and then stabilized, indicating a more stable genomic status [Figure 2].

**Figure 2:**
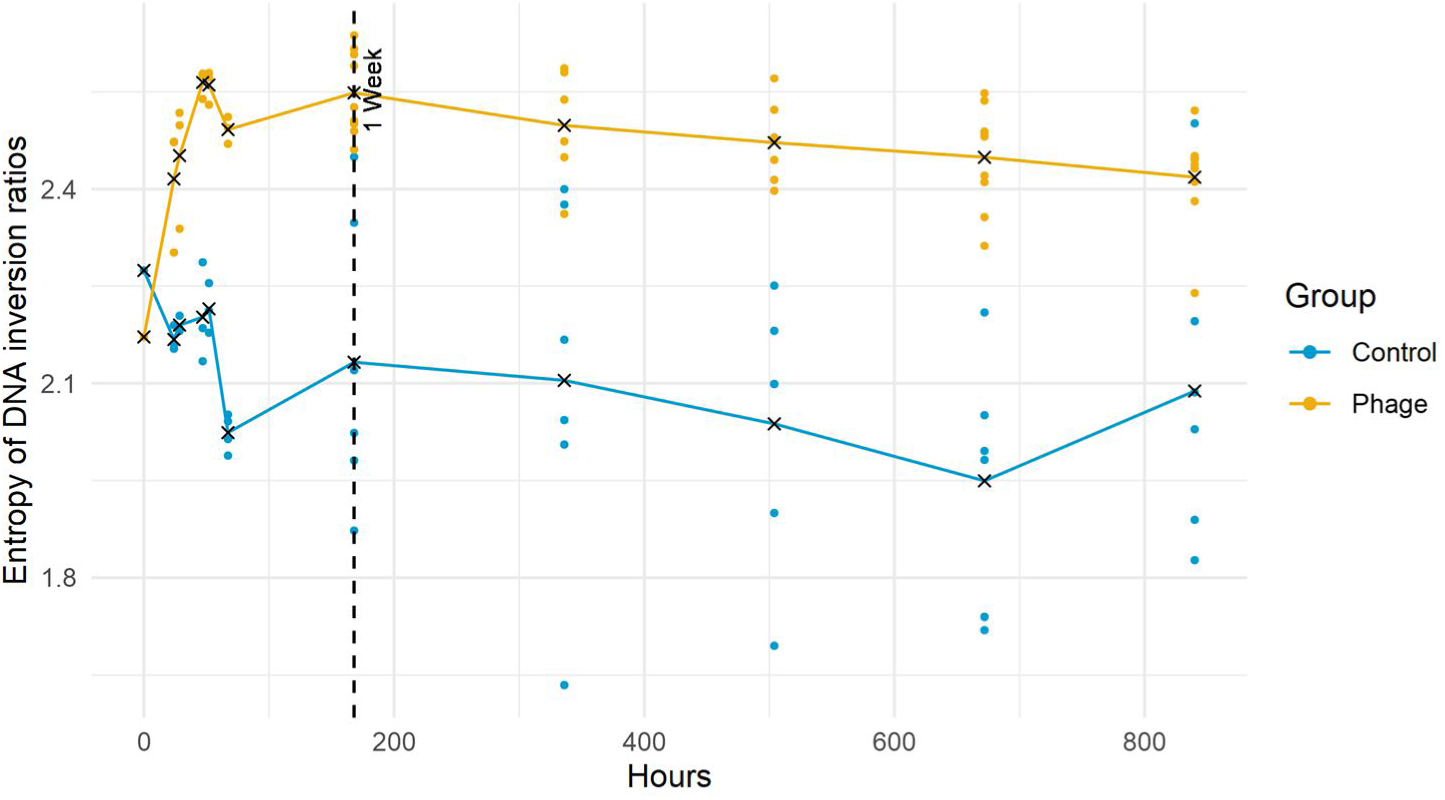
Phage exposure induces entropy changes on overall DNA inversion ratios. Longitudinal entropy analysis on *B. fragilis* DNA inversion ratios from mice monocolonized with *B. fragilis* (CTRL; blue) or colonized with *B. fragilis* and inoculated with Barc2635 (Phage; yellow). The dots represent each fecal sample, the ‘X’ represents the mean entropy. The vertical black dashed line represents the one-week timepoint (168 hours).

This dynamic was particularly pronounced in the PS promoter regions, where the phage-exposed group displayed a rapid and sustained increase in entropy. In contrast, the PVRs (excluding PS promoters) exhibited a subtler increase in entropy, with less pronounced differences between the phage-exposed and control groups [Figure S1].

The introduction of bacteriophage triggered shifts in several invertible regions within the *B. fragilis* genome. Among the seven polysaccharides with invertible promoters, PSG demonstrated the fastest, most dramatic shift among the seven polysaccharides with invertible promoters. Under baseline conditions, PSG typically remained in the “OFF” state with a near 0% “ON” ratio, *in vitro* and *in vivo*. Upon exposure to the bacteriophage, PSG rapidly transitioned to a 75% “ON” orientation within the first 24 hours, followed by a decline to approximately 30% “ON” by 36 hours. In contrast, the control group showed a gradual increase in PSG “ON” orientation to approximately 20% over one week [Figure 3;PSG]. Notably, polysaccharide E (PSE) showed consistent shifts across the two groups, shifting from an initial 55% of the population with the “ON” orientation *in vitro* to 85% under *in vivo* conditions before stabilizing [Figure 3;PSE].

**Figure 3:**
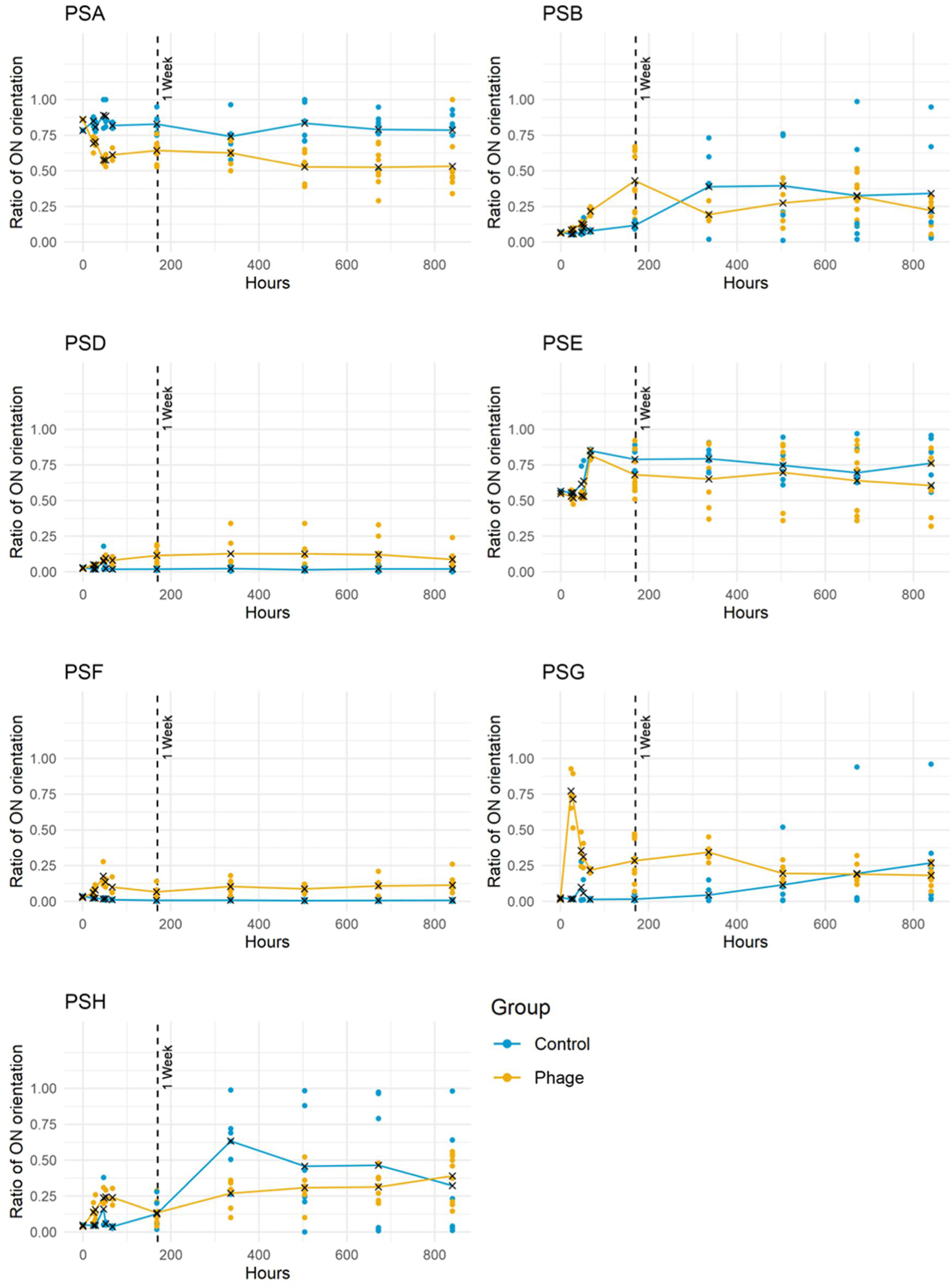
Phage exposure alters the *B. fragilis* polysaccharides’ invertible promoter’s orientations. Ratio of *B. fragilis* polysaccharides’ invertible promoter’s “ON” orientations measured at different hours in fecal samples of mice monocolonized with *B. fragilis* (Control; blue) or colonized with *B. fragilis* + Barc2635 (Phage; yellow). The dots represent each fecal sample, the ‘X’ represents the mean ratio. The vertical black dashed vertical line represents the one-week timepoint (168 hours).

PSA exhibited a response consistent with our previous observations^25^. Its initial “ON” orientation was approximately 75% *in vitro*. Upon exposure to phage, PSA transitioned to an “OFF” state, with the ‘ON’ ratio declining to 50% and stabilizing at this level. The control group maintained a consistent PSA ‘ON’ ratio of approximately 75% throughout the experiment [Figure 3;PSA].

Other polysaccharides’ invertible promoters demonstrated distinct response patterns. PSB and PSH showed accelerated shifts to the “ON” state compared to control conditions [Figure 3;PSB,PSH]. PSF and PSD, which typically remained “OFF” *in vivo*, exhibited a subtle activation in the phage-exposed group, maintaining a low “ON” ratio of approximately 10% [Figure 3;PSF].

Beyond polysaccharides, other phase-variable regions (PVRs) displayed distinct responses to phage exposure. Of particular interest was PVR1, an invertible region located between genes encoding a recombinase (BF9343_2694) and an extracellular polysaccharide (EPS)-related membrane protein (BF9343_2695). In the phage-exposed group, this region shifted from predominantly “OFF” to approximately 80% “ON” orientation, remaining elevated throughout the experiment. The “ON” orientation of this region enhances the expression of downstream EPS, potentially leading to increased bacterial capsule size and decreased access to the bacterial membrane^29^. In contrast, the control group showed minimal activation of this promoter, being mostly “OFF” and stabilizing its “ON” ratio below 12% [Figure 4;PVR1].

**Figure 4:**
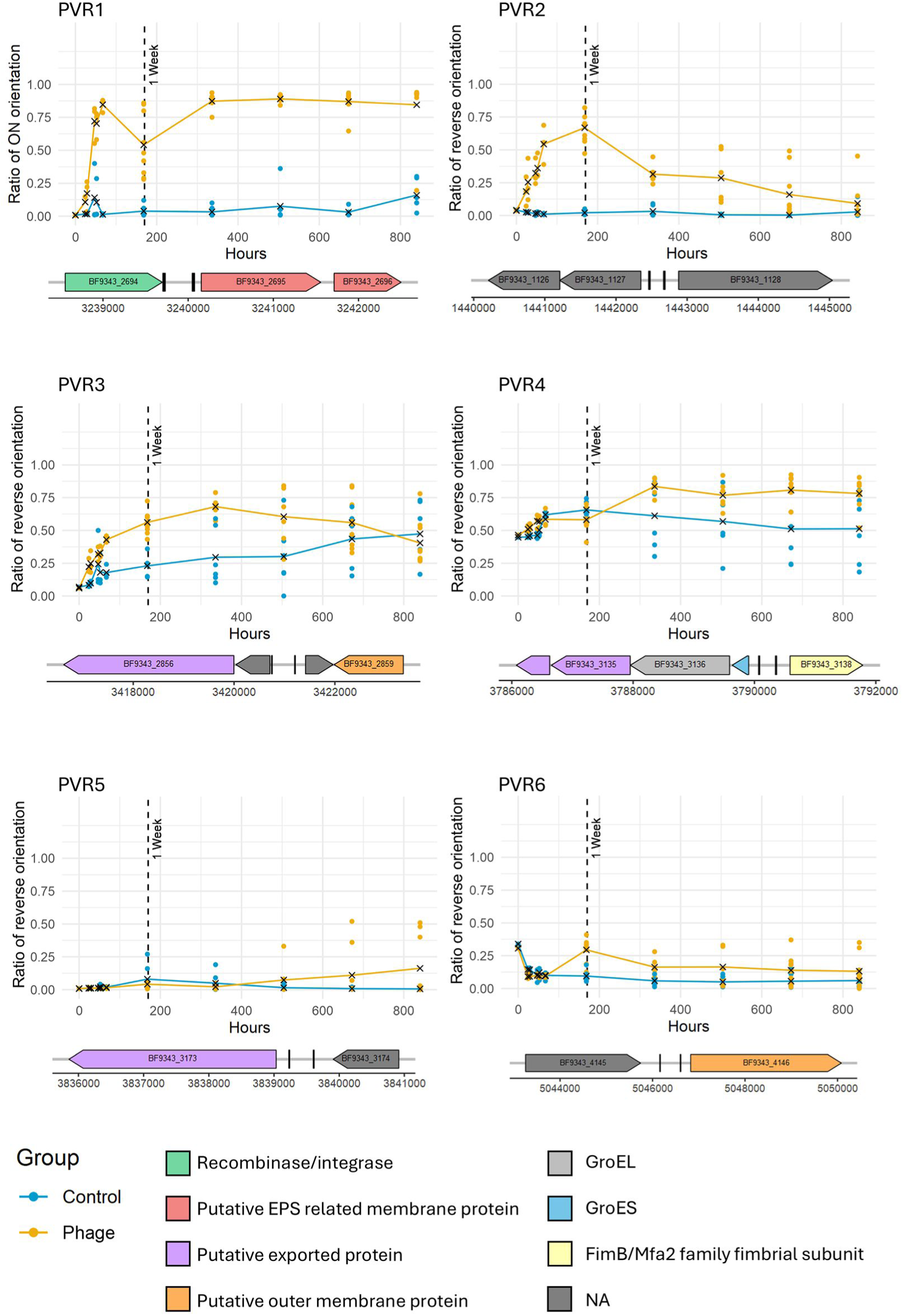
Phage exposure alters the *B. fragilis* invertible promoter’s orientations. Ratio of *B. fragilis* invertible promoter’s orientations measured at different hours in fecal samples of mice monocolonized with *B. fragilis* (Control; blue) or colonized with *B. fragilis* + Barc2635 (Phage; yellow). The dots represent each fecal sample, the ‘X’ represents the mean ratio of orientation. Black dashed vertical line represents the one-week timepoint (168 hours). Genes are represented by colored arrows, and the inverted repeats are represented by black vertical lines in the genomic map accompanying the graph for each invertible region. When applicable, the orientation was designated as “ON” according to the *Bacteroides* promoter motif direction (see methods).

Other invertible regions we surveyed covered a variety of genomic loci, including those associated with outer surface components [Figure 4]. In contrast to the EPS-related region in PVR1, these other invertible sites exhibited more variable responses to phage exposure. PVR2, the invertible region located between genes encoding an ABC transporter substrate-binding protein (BF9343_1127) and a SLBB domain-containing protein (BF9343_1128) showed an increase in the reverse oriented reads after phage exposure. This peaked after one week (mean 66.8%), but then steadily declined back to control levels (lower than 12%) by day 35 [Figure 4;PVR2]. Similarly, PVR3, located near a putative exported protein gene (BF9343_2856) also showed an increase in reverse oriented reads in both the phage-exposed and control groups, but the phage condition demonstrated a more pronounced peak, reaching a mean of 68.13% after two weeks, compared to the control that constantly increased at lower rate and reached a peak of 47.36% on day 35 [Figure 4;PVR3]. Another region, PVR4, located proximal to a gene encoding an outer membrane component (BF9343_3138) of the FimB/Mfa2 family fimbrial subunit, exhibited an increase from *in vitro* (45.6% ratio) to *in vivo* conditions in the control group. The ratio of reversed oriented reads rose to 65.61%, after one week, then slowly reverted to a mean of 51.29% by day 35. Under phage exposure, there was a further increase to 83.49% after two weeks, which then stabilized [Figure 4;PVR4]. In contrast, PVR5 near a putative exported protein gene (BF9343_3173) and PVR6 near a putative outer membrane protein gene (BF9343_4146) maintained relatively low ratios of reverse oriented reads overall. However, PVR5 did show a slight increase towards the end of the experiment, while PVR6 remained consistently higher with phage exposure than the control [Figure 4; PVR5,PVR6]. Notably, the control groups for these invertible regions generally displayed minimal fluctuations in orientation over time, distinct from the more dynamic shifts observed under phage exposure conditions. Additional surveyed PVRs showed relatively low ratios of reverse oriented reads [Figure S2].

We used the PVR ratios to create a principal component analysis (PCA) plot, which allowed us to visualize the differences between the experimental groups and timepoints [Figure 5].

**Figure 5:**
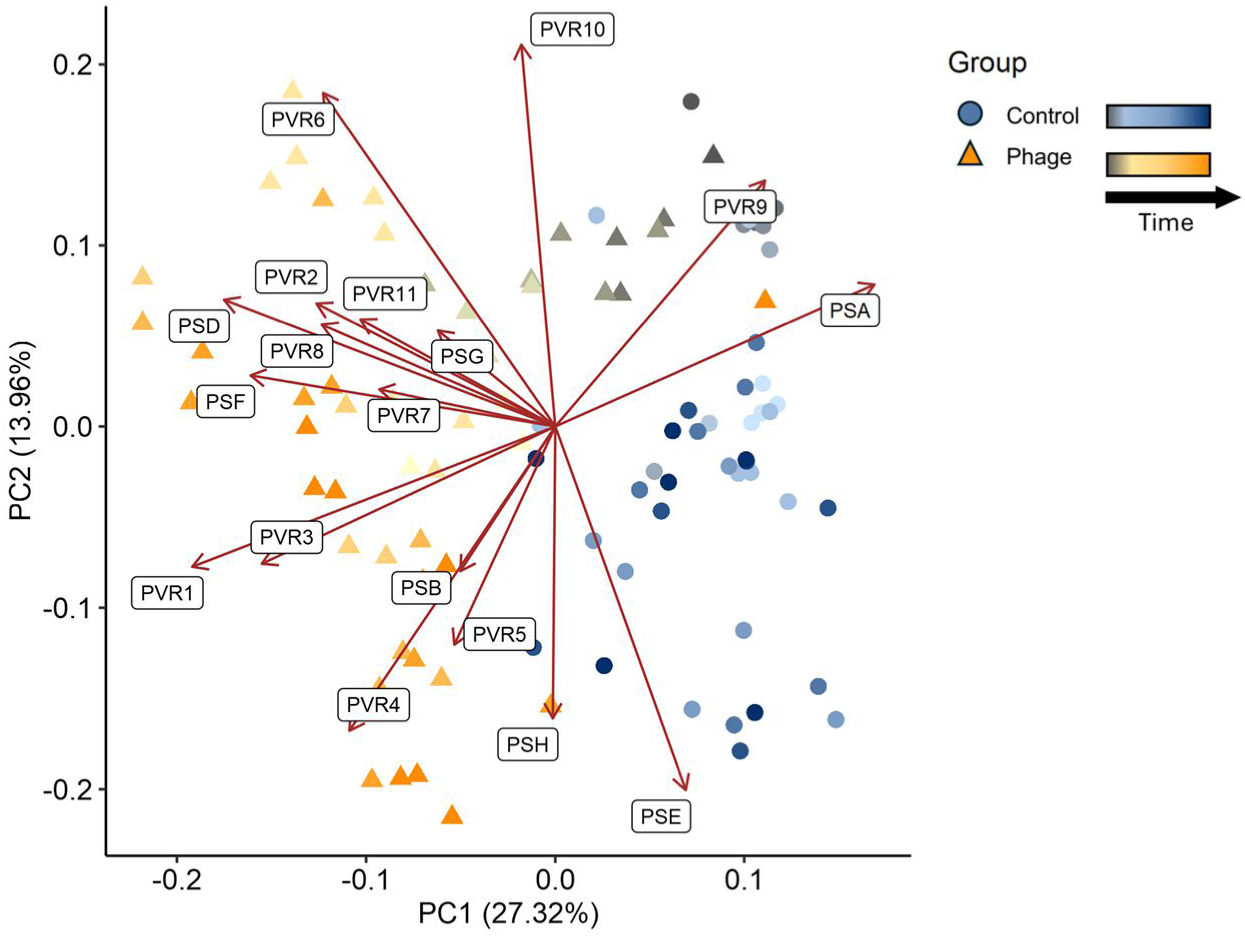
Principal Component Analysis (PCA) of PVRs inversion ratio profiles across experimental conditions and timepoints. Principal components analysis (PCA) of Polysaccharides’ invertible promoters and PVRs inversion ratio profiles of different groups and timepoints. Each point represents a single fecal sample, colored according to group mice monocolonized with *B. fragilis* (Control; blue circle) or colonized with *B. fragilis* + Barc2635 (Phage; yellow triangle), and timepoints – from gray (0 Hours) to darker shades of either group as time progresses. Loadings are represented as red arrows, varying in length according to their contributions.

The starting point samples appeared closely clustered together, indicating similar profiles across the PVRs at the beginning of the experiment. However, as the experiment progressed, the samples began to diverge, revealing distinct trajectories for the control and phage-exposed groups. Along the primary axis, PC1, which explains 27.32% of the variance, we observed a clear separation between the control and phage-exposed samples [Figure 5]. This shift was driven by most of the PVRs measured, with high loading values for PVR1 and PSA (turning “OFF” in the phage-exposed group).

Secondly, the samples also diverged along the secondary axis, PC2, which accounts for 13.96% of the variance [Figure 5]. This separation appears to be largely driven by the temporal dynamics, with samples collected at later timepoints diverging from the earlier timepoints. This shift indicates that DNA inversions in PSB, PSE, PSH, PVR4, and PVR5 might be related to the bacteria adapting to *in vivo* conditions.

### Dynamic changes of B. fragilis Type 1 Restriction-Modification through invertible DNA regions

The T1RM system in *B. fragilis* demonstrated dynamic shifts in specificity gene transcripts combinations over time. The T1RM system consists of a type 1 restriction-modification complex, which includes a restriction enzyme (HsdR, BF9343_1754), a methylase (HsdM, BF9343_1756), and four genes or partial genes (BF9343_1757 through BF9343_1760) that encode six distinct modules (N-terminal or C-terminal halves) of specificity proteins (HsdS)^13^. Only one specificity gene is actively transcribed—the gene located in the expression locus just downstream of *hsdM*. Two half-genes can encode the N-terminal portion of the specificity protein (denoted as ‘Heads’: 57H and 60H, [Figure 6A]) and four half-genes that can invert into the expression locus to comprise the C-terminal portion (denoted as ‘Tails’: 57T-60T, [Figure 6A]). Inversion events between the HsdS half-genes result in eight different combinations of specificity proteins, depending on the DNA orientation [Figure 6A].

**Figure 6:**
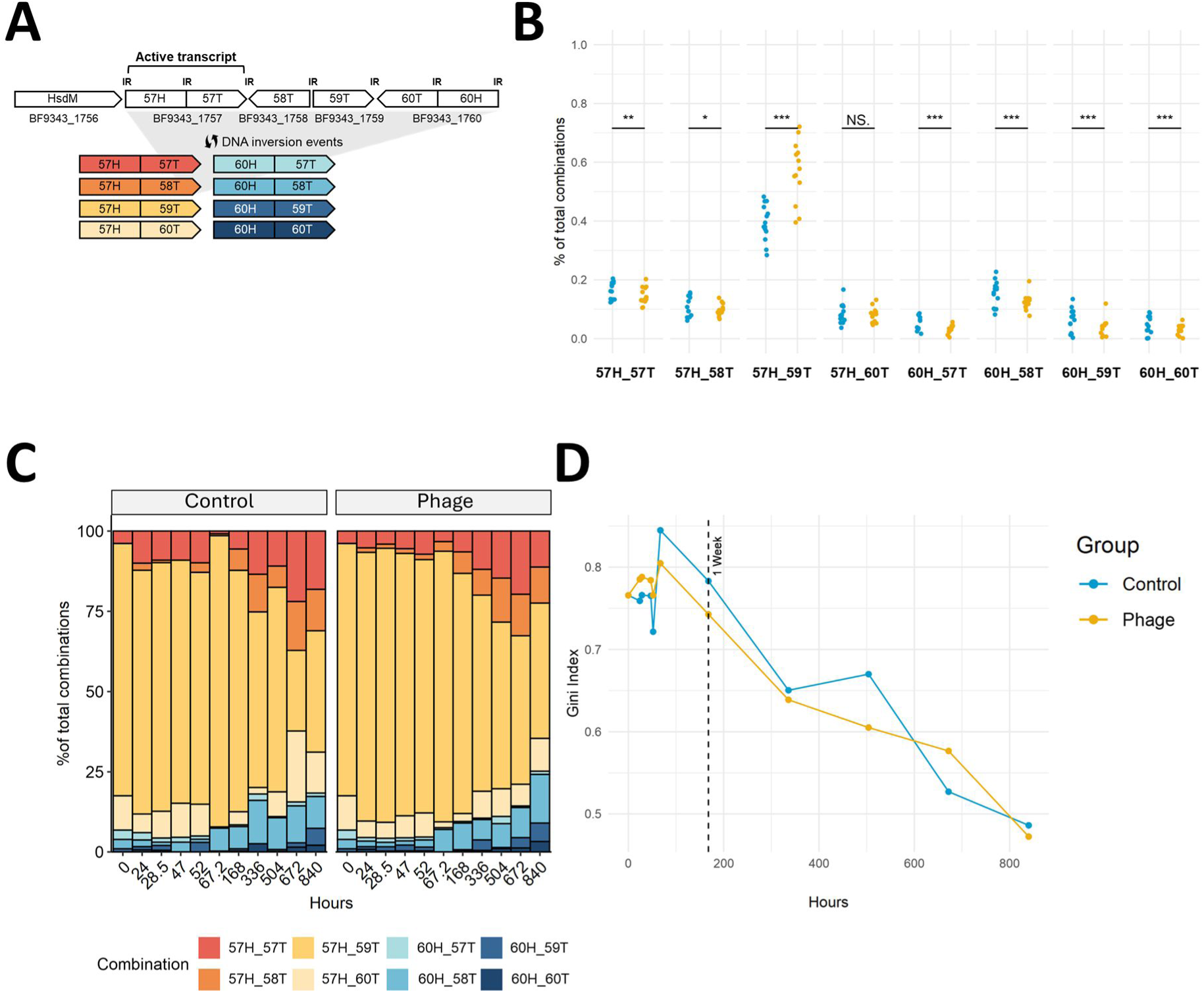
Changes in the distribution of *B. fragilis* Type 1 Restriction-Modification system specificity gene transcripts combinations through time. (A) Gene map of the T1RM region: hsdM-methylase; BF9343_1757-60, represents the phase-variable inverting regions of hsdS (specificity genes) with the locations of inverted repeats (IR) between them. (B) Relative percentages of specificity gene transcripts at the expression locus at day 10, in fecal samples of mice monocolonized with *B. fragilis* (Control; blue) or colonized with *B. fragilis* + Barc2635 (Phage; yellow). The dots represent each fecal sample, \**p* < 0.05, \*\**p* < 0.01, \*\*\**p* < 0.001, NS = not significant (Wilcox rank-sum test). (C) Relative abundances of sequenced reads supporting each of the 8 specificity gene transcripts in fecal samples of mice monocolonized with *B. fragilis* or colonized with *B. fragilis* + Barc2635 through time. (D) Gini index, measuring distribution inequality of the relative abundances of sequenced reads supporting each of the 8 specificity gene transcripts in fecal samples of mice monocolonized with *B. fragilis* (Control; blue) or colonized with *B. fragilis* + Barc2635 (Phage; yellow) through time. Black dashed vertical line represents the one-week timepoint (168 hours). The Gini index indicates how evenly the bacterial combinations are distributed; lower values indicate more equality in distribution.

To determine whether the T1RM system exhibits structural variation in response to phage exposure, we conducted an initial experiment at day 10 post-inoculation, comparing bacteria isolated from phage-exposed mice to control mice. Our results revealed significant differences in multiple combinations [Figure 6B]. The 57H_59T combination showed the highest proportion in both groups, and was enriched in the phage group, while other combinations demonstrated reduced or stable levels.

Specificity proteins of T1RM systems recruit restriction enzymes and methylases to the genomic recognition sites. Each of the eight specificity proteins of the *B. fragilis* T1RM system recognizes a different genomic target sequence^13^. Analysis of the Barc2635 genome revealed two recognition sites of the 57H-59T (GACN5CTG), one on each strand, in locations MN078104.1:29472-29483 and MN078104.1: 35181-35170, and one recognition site for 57H-60T (GACN6TGC), MN078104.1:11867-11856. To note, these specificity proteins require at least two recognition sites, in proximity to each other, in order to function^30–32^.

Based on these findings, we sought to determine whether structural variation in the T1RM system occurred before or after the shifts we saw in genomic orientations of the studied PVRs. At first, the 57H_59T combination predominated in both the phage-exposed and control groups, maintaining consistently high proportions through the first 168 hours (7 days) [Figure 6C]. After this initial period, a notable decrease in the 57H_59T combination emerged, accompanied by an evident expansion in the diversity of T1RM combinations.

This trend towards a more variable distribution of T1RM combinations became increasingly evident after the first week and continued throughout the remainder of the experiment in both groups. To assess the inequality in combination distributions, we calculated the Gini index for each timepoint and group. The Gini index initially showed high values for both groups, indicating the dominance of a single combination (57H_59T). Over time, the Gini index progressively declined, reflecting the emergence of a more even distribution of combinations. Interestingly, after the first 72 hours the phage-exposed group exhibited consistently lower Gini index values compared to the control group [Figure 6D].

## Discussion

Our findings show that bacteria adapt to host colonization and phage exposure through multiple mechanisms that unfold over different timescales. On the population level, the bacteria patterns exhibit oscillations over time, with fluctuations at higher frequency during the first week, suggesting an adaptation period to the mammalian host which stabilizes at about two weeks. Upon phage exposure, the bacterial oscillations fluctuate with lower amplitudes during the first week, suggesting a higher predation. Gradually, the oscillations frequencies are reduced in both groups (with or without phages), however, the oscillations pattern retained throughout time, suggesting a continuous, dynamic, interaction between phage and bacteria. At a later stage, an increase in phage levels accompanied by relatively stable levels of bacteria may reflect the adaptation of the phage to the bacterial changes.

To study the bacterial changes on the functional level (i.e. bacterial functional plasticity) in response to colonizing the mammalian host and to phage exposure, we analyzed bacterial DNA inversions, which can regulate functional alterations, and as a consequence coping with phage existence. In this regard, we find valuable information both in the extent of DNA inversions and in the genes affected by these genomic alterations.

We identified reversible DNA inversions, alluding to bacterial functional plasticity, predominantly in genomic regions controlling outer surface components including secreted proteins, membranal molecular transporters, potential fimbriae, extracellular and outer surface polysaccharides. Since the outer surface of the bacteria is the first physical entity interacting with external factors, such as both the mammalian host, and phage, it is not surprising that these components varied. Indeed, a previous study identified outer surface PSs of *Bacteroides thetaiotaomicron* as targets of its associated bacteriophages. Some PVRs were common in both groups (with and without phage), and therefore predominantly related to host adaptation. Interestingly, the number and diversity of altered PVRs upon exposure to phage were markedly higher, both in number and in diversity. Several outer surface polysaccharides, an ABC transporter, and an outer membrane component of fimbriae, are among the PVRs which altered in bacteria specifically upon exposure to phage. Among the seven phase variable PS promoters of *B. fragilis* that altered in response to phage – PSA was more frequent in its “OFF” orientation, whereas PSG, PSF, and PSD in their “ON” orientations. The PS of *B. fragilis* exhibit a hierarchical regulation, where the PSs loci include trans locus inhibitors (UpxZ), each able to silence a different repertoire of PSs^33^. For example, the PSF trans locus inhibitor, UpfZ, inhibits seven PSs, UpgZ inhibits six PSs, both inhibiting PSA, while UpaZ, UpbZ and UphZ inhibit five PSs, and the rest inhibit two PSs or less.

The chronological sequence of these adaptive changes provides insights into potential cause-and-effect relationships, despite our limited understanding of the drivers of DNA inversion. For example, the PSG promoter exhibited substantial changes within the first 24 hours - starting with a sharp increase in frequencies of the “ON” orientation and followed by a sharp drop to a more frequent “OFF” orientation in the population. Since PSG inhibits multiple PSs^34^, we hypothesize that this initial increase in the “ON” orientation results in a substantial reduction of expressed polysaccharides, possibly veiling from phage^14^. On the other hand, the subsequent “OFF” orientation of the PSG promoter enables the expression of multiple PSs, possibly promoting the phage-host dynamic “arms-race” [Figure S1]. Both strategies can be explained as response to stress: the first quickly shutting off molecules with potential detrimental effects (e.g. PS can provide molecular attachment and entry-points for the phage^35^), and the latter, inducing the expression of multiple phenotypes increasing the probability of the bacterial population to survive by applying a bet-hedging strategy, known to be advantageous in stress conditions^36^. Another molecular system we found relevant to focus on, is the type 1 restriction-modification (R-M) system. T1RMs are systems that potentially can provide protection against bacteriophages, by identifying and restricting foreign DNA. They are composed of specificity proteins, which recognize specific “recognition sequences” in the DNA, and recruit either methylases, for DNA methylation, or restriction enzymes, to restrict the DNA, recognized as foreign. We previously characterized a phase variable T1RM system in *B. fragilis* and showed that DNA inversions of this system lead to altered global transcriptional programs, including controlled expression of different capsular polysaccharides. In this study, we find DNA inversions of this T1RM system over time, both upon adaptation to the host and in response to phage exposure. Upon phage exposure, the dominated specificity gene of this phase-variable T1RM system is the only specificity protein (out of eight) which can recognize the DNA of Barc2635 bacteriophage and potentially restrict it, preventing phage infection. ^30–32^. Intriguingly, the transcriptional program controlled by this T1RM orientation upregulates the PSF biosynthesis locus^13^, which, in turn, downregulates the expression of PSA^33^. Consistently, and in parallel, we find an increase in the ‘OFF’ orientation of the PSA promoter, over time, which is an independent DNA inversion event, regulated by a dedicated invertase^18^. These bacteriophage-induced phase variable alterations in *B. fragilis* prolonged for over 4 weeks (i.e PSA ‘OFF’, PSF ‘ON’). This indicates high demand in shutting off PSA during exposure to phage, with multiple mechanisms regulating this outcome.

An open question, to date, is whether bacteria respond to stress by increasing the diversity of their molecular repertoire, or vice *versa*. On one hand, increasing diversity of the expressed molecules provides the bacteria with more functional capabilities, enabling bet-hedging^5^, and on the other hand, such increased variability might require excess energy and complex bacterial economy (in regard to maintaining molecular diversity). Our results suggest that upon encountering both a new host environment and phage stress, the bacterial population shows an increase in their molecular diversity, measured by higher entropy and a relatively large number of genomic cites being inverted. Interestingly, the DNA inversions of the PSs were rapidly altered in both groups (with and without phage), however, at a much higher rate and to a much higher degree in the group exposed to phage, with both groups displaying higher entropy compared to the original inoculated population.

When diving into all measured DNA inversions, excluding the PSs, the two groups display a distinct pattern of entropy, where the entropy of the control group is gradually reduced, and both groups maintain their level of entropy over time.

In contrast to the PS inversions, the T1RM system, despite its critical role in cleaving foreign DNA, including phage, demonstrated a much slower adaptation pattern throughout the experiment. Albeit having one specificity gene combination favorable for phage DNA restriction, which indeed was more represented in the phage group, the T1RM system in both control and phage groups, altered in comparison to the initial inoculation, and gradually became more evenly dispersed, with all eight orientations represented. We hypothesize that this higher diversity, achieved over time, reflects a bet-hedging strategy in response to the fluctuating conditions. Phage can acquire methylation patterns upon encountering their host and its methyltransferases in order to resist host restriction^37^. The observed spread between the eight T1RM combinations, over time, might be explained by phage-acquired methylation patterns, resulting in phase-variable T1RM diversity, since the bacteria require multiple T1RM orientations to cope with the phage.

Our previous study showed that the PSA promoter is more frequently in its “OFF” orientation in IBD patients, concomitant with high abundance of *B. fragilis* associated gut bacteriophages. Further assessment indicated no phage preference for the presence of the PSA molecule on the bacterial membrane. This study also reveals a higher frequency of the “OFF” orientation of the PSA promoter upon phage exposure, which prolongs throughout the experiment. In this study, we expand our findings showing that the PSA promoter “OFF” orientation is accompanied by an “ON” orientation of other, related PSs. This emphasizes the importance of outer surface PSs variability in the bacteria population during stress conditions, aligned with the observed variability in additional phase-variable genomic regions.

Intriguingly, PSA is an outer surface immunomodulatory molecule, inducing regulatory T cells (Tregs) and the interleukin (IL)-10 cytokine, rendering anti-inflammatory effects on the host^19^. In our previous study^25^, where the PSA promoter orientation was oriented “OFF” in one timepoint, we confirmed its translation to a reduction in colonic Tregs. This current study indicates a prolonged phage associated “OFF” orientation of the PSA promoter, which persists for the whole duration of the experiment (five weeks). Notably, this study reveals multiple DNA inversions in outer surface molecules, in addition to PSA, and therefore merits future studies on their potential effects on the host.

A recent study used primarily transcriptomics to characterize the mechanisms that allow phage-bacteria co-existence in different bacteria (other *Bacteroides* and a *Parabacteroides* species) cultured *in vitro*^38^. The researchers found transcriptional alterations in capsular polysaccharides (CPS) as a primary mechanism supporting phage co-existence in these bacteria, and that additional resistance strategies, such as metabolic adaptations and regulation of surface proteins, are also activated to ensure bacterial survival^38^.

Our study complements these findings both by providing an *in vivo* perspective on *Bacteroides* adaptation to phage exposure, and by emphasizing the role of genomic DNA inversions, in mediating these adaptations. By analyzing both short- and long-term dynamics, we reveal how these genomic changes drive bacterial responses to phage predation and to the host gut environment over time. Together, these two studies offer a comprehensive understanding of phage-bacteria interactions, from molecular mechanisms to their ecological relevance.

Understanding the clinical implications of bacterial adaptation to phages is crucial for advancing microbiome-based therapies. The virome plays a significant role in shaping health and disease outcomes^39–41^. Fecal microbiota transplantation (FMT), which transfers a complex community of both bacteria and viruses, has shown effectiveness in treating a variety of diseases, highlighting the potential of microbiome-based interventions^42,43^. Our study demonstrates that phage exposure can lead to DNA inversions in bacteria in multiple genomic regions, potentially altering their functionality. This may provide mechanistic explanations for the effectiveness of fecal virome transplantation (FVT)^44–46^, as its therapeutic benefits could arise not only from shifts in bacterial composition but also from phage-driven changes in bacterial functionality.

Our study highlights the central role of DNA inversions in the adaptation of *B. fragilis* to the mammalian gut and to bacteriophage exposure. Although the phage is lytic, the bacterial population persisted over time. The observed DNA inversions, primarily in outer surface molecules and polysaccharides act as rapid and flexible strategies, enabling the bacteria to respond to external pressures and potentially alter their effects on the host. This dynamic and ongoing process of genomic restructuring was marked by variability in both the PVRs and the T1RM system inversion patterns. The shift towards a more even distribution of specificity gene combinations within the T1RM system, alongside the increased entropy of DNA inversions throughout the genome, suggests a bacterial bet-hedging strategy.

Our results underscore the importance of considering temporal factors in host-microbe interaction studies. Bacterial adaptation to the host environment, even in the absence of phage pressure, appears to occur both in an initial phase within days and a second phase emerging in later stages. This observation might have consequences on microbe-host interactions throughout the course of such adaptations and shall be considered when designing experiments integrating optimal timing of analysis. This study advances our understanding of bacterial adaptation mechanisms and their temporal dynamics, while also offering important insights on microbe-host interactions. Additionally, it emphasizes the importance of high temporal resolution, especially during early adaptation phases. Future research could focus on elucidating the molecular drivers of DNA inversions and validating these findings in human microbiome contexts.

## Limitations

Several experimental considerations warrant discussion. This study was limited to examining the interaction between a single phage and a single bacterial species. As such, the findings may not directly translate to other bacterial species or more diverse microbial communities. Additionally, since we focused on DNA inversions as a mechanism of genetic plasticity and adaptation, by analyzing changes at the genome level of the bacteria, further research is needed to explore the effects of other genetic mechanisms that may influence these dynamics and their impact on gene expression. Moreover, this study centered on bacterial adaptation to the phage without investigating the reciprocal phage adaptation. Regarding the time scale of our sampling, at later stages of our experiments – we sampled on a weekly basis, we may have missed finer-scale dynamics and transient changes that occur between sampling points. Finally, our experimental model utilized monocolonized mice, which limits its generalizability to more complex *in vivo* systems, such as specific pathogen-free (SPF) environments, defined microbial communities, or other multi-microbial contexts. With these limitations notwithstanding, our study provides valuable insights into the dynamics of phage-host interactions and the significance of studying bacterial genomic structural variation as a mechanism of adaptation.

## Availability of data and materials

Data of metagenomic and long-reads sequencing have been deposited at the SRA database under project accession numbers PRJNA1199331 (PVRs and Type 1 RM system specificity gene transcripts combinations of longitudinal experiment) and PRJNA948162 (Type 1 RM system specificity gene transcripts combinations at day 10 after phage exposure).

## Author contributions

S.C., H.H., R.K.-D., D.K.-K., T.G., and N.G.-Z. conceived and designed the project. S.C., R.K.-D., J.Z., H.H., D.K.-K. performed and analyzed the experiments. S.C., R.K.-D., J.Z., H.H., T.G., and N.G.-Z. wrote the manuscript. S.C., and R.K.-D. contributed equally to the study. All authors read and approved the manuscript. N.G.-Z. conceived and planned the study, supervised it, interpreted the experiments, and wrote the manuscript.

## Acknowledgments

We would like to thank the Geva-Zatorsky lab members for fruitful discussions and contributions throughout this study. Special thanks to Drs. Svetlana Fridman and Nadav Ben-Assa for conductive consultations. We thank Prof. Laurie Comstock for enabling us to seed this research directions, as well as for productive and valuable discussions and guidance. We thank Prof. Juan Joffre for isolating the Barc2635 bacteriophage. BiotaX labs for metagenomics sequencing; Dr. Neeru Bhardwaj and Mrs. Margalit Beck Bobritsky for help in sampling, as well as Mr. Shimon Neriah Michaeli & Ms. Efrat Oded for DNA extraction.

## Material and Methods

### *B. fragilis* and bacteriophage Barc2635 culture conditions

*Bacteroides fragilis* NCTC 9343 (RefSeq: NC_003228.3) was obtained from ATCC and stocks in 25% glycerol were kept at −80°C. Bacteria were thawed on BHIS plates for 2-5 days in an anaerobic chamber then isolated colonies were obtained and transferred to 4ml liquid BHIS tubes overnight (o/n) at 37° under anaerobic conditions.

Bacteriophage Barc2635 (GenBank: MN078104) was isolated from sewage as previously described (Carasso et al.^25^). Prior to experiments, phage propagation was performed as follows: individual, well-isolated plaques were picked with a sterile needle and transferred to 5 mL BRPM broth. A 1 mL aliquot of *B. fragilis* NCTC 9343 culture in exponential growth (OD 0.5–0.8) was added to the broth, and the mixture was incubated anaerobically at 37°C for 18 hours. Following incubation, the culture was treated with chloroform (1:10 v/v), vigorously mixed for 5 minutes, and centrifuged at 16,000 × g for 5 minutes. The resulting supernatant containing the phage was filtered through a low-protein-binding 0.22 µm pore size polyethersulfone (PES) membrane filter (Millex-GP, Millipore, Bedford, MA) to remove bacterial debris.

Phage stocks were stored at 4°C, and the plaque-forming units (PFU) were quantified prior to initiating experiments to ensure consistent phage concentrations.

### Phage-bacteria *in vivo* co-culture experiment

All animal experiments were conducted in accordance with protocols approved by the Institutional Animal Care and Use Committee (IACUC) under approval number IL-105-06-21. Female germ-free (GF) C57BL/6 or Swiss Webster mice, aged 4–5 weeks, were obtained from the Technion GF colony. Mice were housed in a germ-free care facility, maintained on a 12:12-hour light-dark cycle at room temperature, and provided with food and water *ad libitum*.

Experimental groups included mice administered with *B. fragilis* NCTC 9343 alone or in combination with bacteriophage Barc2635 (n=3-6 per group, in three experiments). All administrations were performed via oral gavage. For the *B. fragilis* only group, overnight cultures of bacteria were directly used (100 µL per mouse). For the phage-exposed group, bacteriophages were mixed with the overnight *B. fragilis* culture at a ratio of 10 phages to 1 bacterium immediately before gavage (100 µL per mouse).

Mice were monitored throughout the experiment and were sacrificed either at the designated endpoint or earlier in cases of contamination.

Fecal samples were collected at predetermined timepoints:

0 hr, 4.5 hr, 23 hr, 28 hr, 47 hr, 53 hr, 72 hr, 118.5 hr, 142 hr, 144 hr, 168 hr (1 week), 190 hr, 238 hr, 312 hr, 334 hr or 336 hr (2 weeks), 382 hr, 480 hr, 504 hr (3 weeks), 648 hr, 672 hr (4 weeks), 816 hr, 840 hr (5 weeks)

### Enumeration of Bacteria and Phage

#### Bacteria Enumeration

Fecal pellets were weighed, suspended, and homogenized in 1,000 µL of sterile PBS. Serial dilutions were prepared in PBS, and the diluted supernatants were plated. Plates were incubated under aerobic conditions to assess contamination and under anaerobic conditions at 37°C for 1–2 days until colonies were countable. Colony counts were multiplied by 100 to account for the total volume, then divided by the fecal sample weight to obtain colony-forming units (CFU) per gram of feces.

#### Phage Enumeration (Plaque Assay)

Filtered diluted supernatants from fecal suspensions were passed through 0.22 µm pore size polyethersulfone (PES) membrane filters with low protein binding. The double-agar layer technique was employed^47^, wherein *B. fragilis* grown to an OD of 0.4–0.7 in BPRM was added to 40°C soft BPRM agar at a 1:10 ratio and plated on BHIS plates. Diluted filtered supernatants were then applied to the plates, which were incubated anaerobically for 1 day. After incubation, bacterial growth covered the plates, and visible plaques were counted. Plaque counts were multiplied and adjusted to determine plaque-forming units (PFU) per gram of feces.

#### DNA Extraction

Fecal samples, collected and stored at −80°C following CFU and PFU enumeration, were processed for DNA extraction using the ZymoBIOMICS DNA Miniprep Kit (Zymo Research) following the manufacturer’s protocol.

#### Illumina Library Preparation and Sequencing

Extracted DNA quality and concentration were assessed using Qubit fluorescence analysis (ThermoFisher, Cat. Q32850). Libraries were prepared using the Illumina Tagmentation DNA Prep protocol with a starting mass of 50 ng DNA and eight cycles of PCR enrichment. Fragmentation yielded DNA fragments averaging 550 bp.

Indexing was performed using IDT for Illumina DNA/RNA UD indexes and Nextera DNA CD indexes (Illumina IDT, Cat. 20027213; Illumina Nextera, Cat. 20018708). The libraries were diluted to 15 pM and pooled in 96-plex. Validation was conducted with 100-cycle paired-end MiSeq V2 runs (Illumina, Cat. MS-102-2002).

Final sequencing was carried out on a NovaSeq 6000 system (Illumina, Cat. 20028312) in S4 mode, with 300-cycle paired-end reads and an average sequencing depth of 30 Gbp per sample. Libraries were loaded at a concentration of 600 pM, incorporating a 1% PhiX control library V3 spike-in (Illumina, Cat. FC-110-3001).

#### MinION Library Preparation and Sequencing

Regions of interest were identified through whole-genome sequencing of samples from the long-term experiment, aligned to the *B. fragilis* NCTC 9343 reference genome (RefSeq: NC_003228.3). Twelve regions (500–800 bp) showing significant phase variation ratio changes between day 7 and day 35 were selected.

Primers targeting these regions were designed using Primer BLAST^48^. Additional primers targeting the seven invertible polysaccharide promoters and the T1RM system specificity gene transcripts combinations were adapted from Lan et al.^49^ and Ben-Assa et al.^13^, respectively. Adapter sequences for MinION barcoding were appended to these primers. A full primer list is provided in the supplementary materials [Table S1].

PCR amplification of targeted regions was performed, with T1RM amplified separately due to its larger size. DNA concentrations were normalized using NanoDrop before barcoding individual samples with the PCR Barcoding Kit (SQK-PBK004, Oxford Nanopore Technologies). Barcoded samples were pooled, and libraries were prepared following the manufacturer’s protocol. Sequencing was conducted on the MinION device, controlled by MinKNOW software (v24.02.16), using MinION flow cells (FLO-MIN114, Oxford Nanopore Technologies).

#### Bacterial genome DNA inversions analyses

The genome of *B.fragilis* NCTC 9343 to create a reference for potential invertible sites. PhaseFinder^11^ v1.0 was used to identify invertible sites in the metagenomics samples. The default parameters of PhaseFinder were used. Results were filtered by removing identified sites with <20 reads supporting either the forward or reverse orientations combined from the paired-end method, mean Pe_ratio <1% across all samples, and sites within coding regions of rRNA products. On these results, the orientation ratio between the B. *fragilis-only* group and phage-exposed group were compared in different timepoints (Wilcoxon rank-sum test).

For the minion sequences, DNA inversion sites detected to vary between groups in the metagenomics sequencing results were used to create a reference of both orientations for every studied region. EPI2ME desktop agent v3.7.3 (Oxford Nanopore Technologies) with the fastq custom alignment analysis was used with said reference.

For phase variable promoter regions, when applicable, “ON”\”OFF” assignment was decided according to the Bacteroides conserved promoter motif (TTG-AT-rich region (19-21bp)-TANNTTTG)^11^

#### Type 1RM combinations analysis

Adapters and barcodes sequences were removed from the reads using Porechop (v0.2.4, available from https://github.com/rrwick/Porechop). Reads were oriented using the ‘Preparing Reads for Stranded Mapping’ protocol (Eccles, D.A. (2019). Protocols.io^50^). The reads were aligned to the PCR forward primer using LASTAL (v.1060)**71**, and then reverse-complemented the reverse-oriented reads. The reads were combined to an all forward oriented file and cropped to the first 1300 bases using Trimmomatic (v.0.39)^51^. The reads were then split according to their alignment to the 1757H-57T or 1757H-60T 5” half sequences using LASTAL. Reads were mapped to the full sequences with Minimap2 (v.2.17-r941)^52^ using the –for-only and asm20 options. Mapped read counts were extracted from the Minimap2 SAM output using SAMtools (v.1.7)^53^.

#### Statistical analysis and plotting

Data was analyzed using non-parametric methods to assess differences between groups and over time. Specifically, pairwise comparisons were conducted using the Wilcoxon rank-sum test for between-group differences at specific timepoints. Principal Component Analysis (PCA) was performed on the ratios of reverse-oriented reads for the identified invertible regions (PS loci and PVRs) to visualize variation across experimental groups and timepoints, using the ‘prcomp’ function in R with centered and scaled data. All statistical analyses were performed using R v4.3.1, and statistical significance was defined as p < 0.05. Plots were generated using the ggplot2 v3.4.3^54^, and the ggpubr v0.6.0^55^ packages in R, with distinct color palettes from the metbrewer package.

## Supplementary materials

**Supplemental Figure S1:**
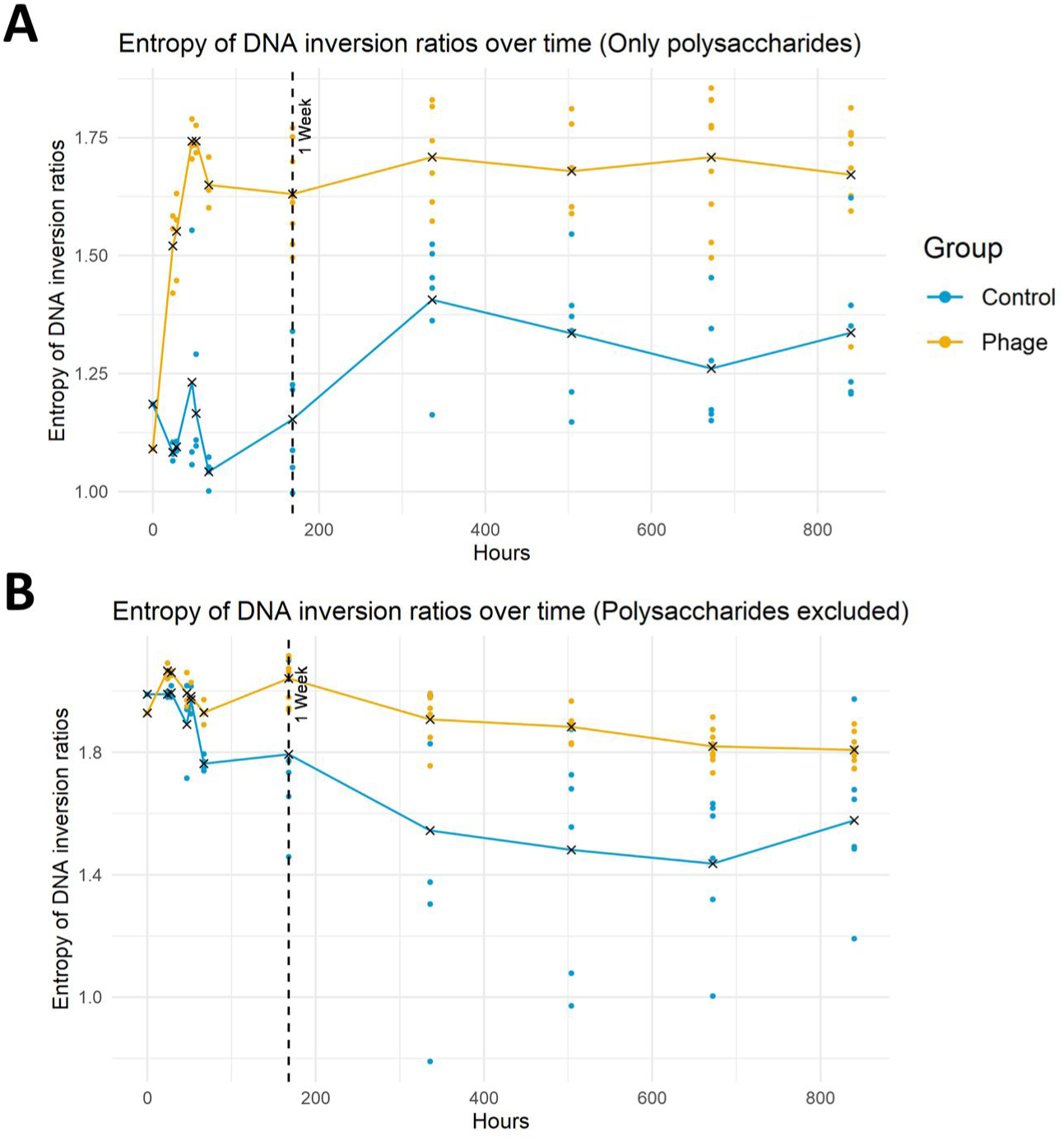
Phage exposure induces entropy changes on DNA inversion ratios. Longitudinal entropy analysis on *B. fragilis* DNA inversion ratios of invertible PSs promoters (A) or PVRs only (B) from mice monocolonized with *B. fragilis* (CTRL; blue) or colonized with *B. fragilis* and inoculated with Barc2635 (Phage; yellow). The dots represent each fecal sample, the ‘X’ represents the mean entropy. Black dashed vertical line represents the one-week timepoint (168 hours).

**Supplemental Figure S2:**
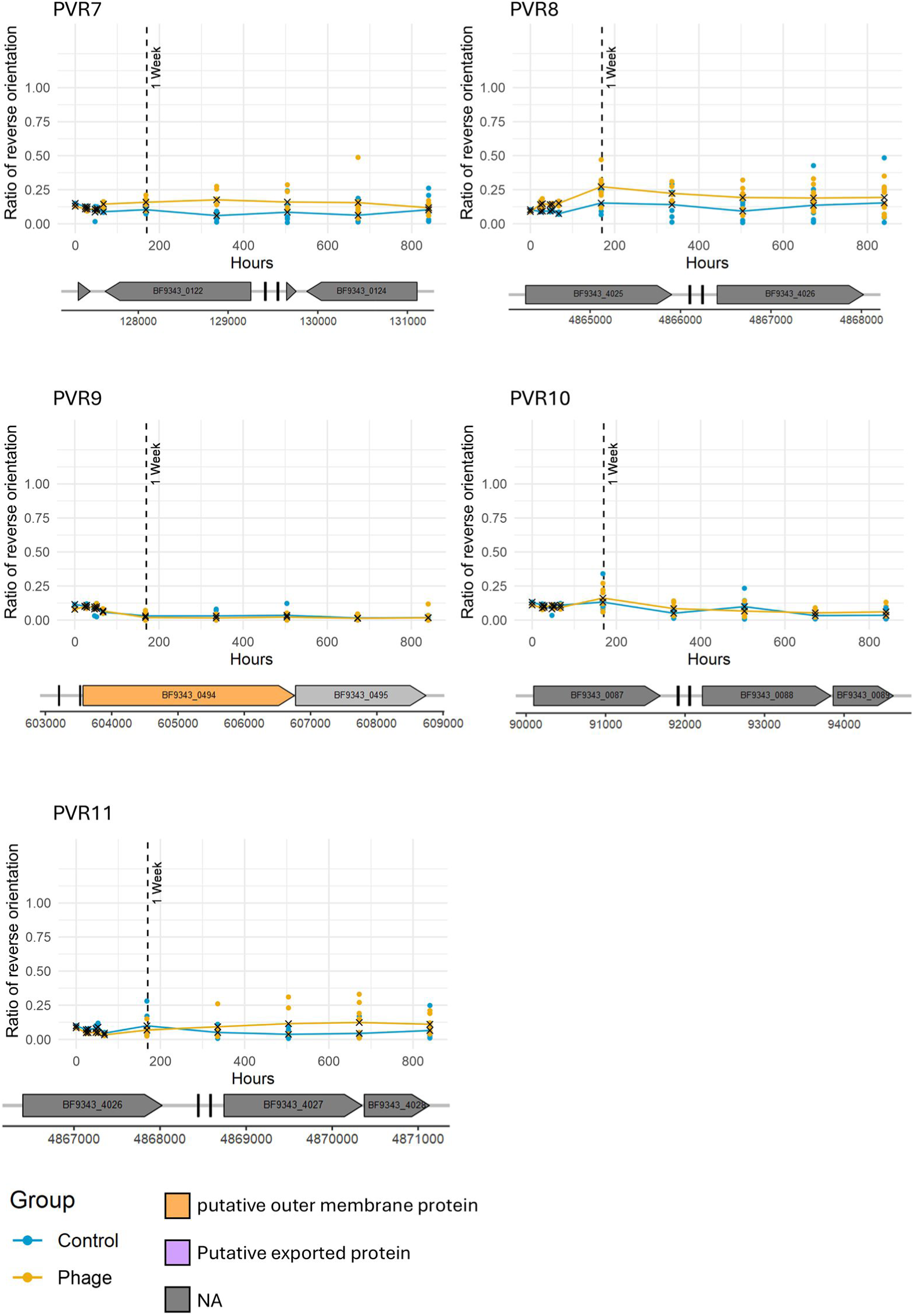
Phage exposure alters the *B. fragilis* invertible promoter’s orientations. Ratio of *B. fragilis* invertible promoter’s “ON” orientations measured on different hours in fecal samples of mice monocolonized with *B. fragilis* (Control; blue) or colonized with *B. fragilis* + Barc2635 (Phage; yellow). ‘X’ represents the mean. Dashed vertical line represents the one-week timepoint (168 hours). Genes are represented by colored arrows, and the inverted repeats are represented by black vertical lines in the genomic map accompanying the graph for each invertible region.

